# Microbiability of meat quality and carcass composition traits in swine

**DOI:** 10.1101/833731

**Authors:** Piush Khanal, Christian Maltecca, Clint Schwab, Justin Fix, Francesco Tiezzi

## Abstract

The impact of gut microbiome composition was investigated at different stages of production (Wean, Mid-test, and Off-test) on meat quality and carcass composition traits of 1,123 three-way-crossbred pigs. Data were analyzed using linear mixed models which included the fixed effects of dam line, contemporary group and gender as well as the random effects of pen, animal and microbiome information at different stages. The contribution of the microbiome to all traits was prominent although it varied over time, increasing from weaning to Off-test for most traits. Microbiability estimates of carcass composition traits were greater compared to meat quality traits. Adding microbiome information did not affect the estimates of genomic heritability of meat quality traits but affected the estimates of carcass composition traits. High microbial correlations were found among different traits, particularly with traits related to fat deposition with decrease in genomic correlation up to 20% for loin weight and belly weight. Decrease in genomic heritabilities and genomic correlations with the inclusion of microbiome information suggested that genomic correlation was partially contributed by genetic similarity of microbiome composition.

## Introduction

The mammalian gastrointestinal tract is a home of a diverse microbiota population which serve various biological functions of the host (Frese et al., 2015). Gut microbiota has recently been the target of many research efforts resulting from the rapid development in molecular technologies and led to a vast influx of “omics” studies (Guevarra et al., 2019). The importance of gut microbiota is widely accepted (Kim et al., 2011), with commensal bacteria often being called the “forgotten organ” of the host (O’Hara and Shanahan, 2006), impacting hosts in a multitude of ways. For example, microbial composition helps in promoting the gastrointestinal health through metabolites, postnatal development, degradation of short chain fatty acids and stimulation of immune system (Mann et al., 2014; Pedersen et al., 2013; Stappenbeck & Herbert, 2016).

Gut microbiome constitutes a portion of the whole genome (Sommer & Bäckhed, 2013; Xiao et al., 2016) and has the potential to affect numerous biological activities that the hosts lack (Pajarillo et al., 2014). Different researchers reported that microbiome has considerable effect on human health and traits (Clemente et al., 2012; Dave et al., 2012; Huttenhower et al., 2012). For example, differences in bacterial species diversity and gene counts between lean and obese individuals have been found (Le Chatelier et al., 2013). The microbial diversity of intestine accounted for significant amount of phenotypic variation for any trait in human and should be accounted when assessing the heritability not only in human but also in plants and livestock (Sandoval-Motta et al., 2017). In livestock, Difford et al. (2016) termed “microbiability” the proportion of total variance explained by microbiome for performance traits of dairy cattle. Difford et al. (2018) reported the effect of microbiota variation in methane production in dairy cows while, Mach et al. (2015) reported the impact of gut microbiome at early life on phenotypes of pig. Gut microbiome also has a significant impact on porcine fatness (He et al., 2016). Camarinha-Silva et al. (2017) reported the presence of a significant effect of microbial composition on daily gain, feed intake and feed conversion rate in swine. Until recently, selection of different traits in pigs has been done with the use of pedigree and genomic information, yet the advantage of incorporating microbial information in the genetic evaluation processes has not been assessed. Few studies have described the relationship of microbial diversity and host e.g. (Guevarra et al., 2019; McCormack et al., 2018), however these were mostly from a nutritional perspective.

Specifically, the contribution of microbial composition to the phenotypic variation of meat quality and carcass composition traits in pigs has yet to be explored and no studies to date have been conducted on the effect of microbial composition at different stages of production on growth and carcass composition. Therefore, the objectives of this study are to estimate the microbiabilities for different meat quality and carcass composition traits; to investigate the impact of intestinal microbiome on heritability estimates; to estimate the correlation between microbial diversity and meat quality and carcass composition traits; and to estimate the microbial correlation between the meat quality and carcass composition traits in a commercial swine population.

## Materials and methods

Animal welfare approval was not needed for this study since all data came from animals raised in a commercial setting by The Maschhoffs, LLC (Carlyle, IL, USA). All pigs were harvested in commercial facilities under the supervision of USDA Food Safety and Inspection Service.

### Animals and sample collection

Data were collected from crossbred individuals that were obtained from 28 funding Duroc sires and 747 commercial F_1_ sows composed of Yorkshire × Landrace or Landrace × Yorkshire. The pigs were weaned at 18.64 ± 1.09 days old and were moved to nursery-finishing facility. Pigs were kept in 334 single-sire single-sex pens with 20 pigs per pen. The test period began the day that pigs were moved to the nursery-finishing facility. During the nursery, growth and finishing period all pigs were fed a standard pelleted feed based on sex and live weight. Details of diet and their nutritional values are provided (see additional File 1). The pigs received a standard vaccination and medication routine. (see additional File 2). End of test (**Off-test**) was reached when the average weight of pigs of each pen reached 138 kg. The average age at off-test was 196.4 ± 7.80 days. Fecal samples for 16S rRNA sequencing were collected as follow. Rectal swabs were collected from all pigs at three stages: weaning (**Wean**), 15 weeks post weaning (**Mid-test**; average 118.2 ± 1.18 days), and off-test. Four pigs from each pen were selected as detailed by Wilson et al. (2016) and their rectal swabs were used for subsequent microbial sequencing. There were 1,205, 1,295 and 1,273 samples for weaning, Mid-test and Off-test respectively. Distribution of samples across families, time points and sex are provided (see additional file 3).

### Illumina amplicon sequencing

DNA extraction, purification, illumina library preparation and sequencing were done as described by Lu et al. (2018). Briefly, total DNA (gDNA) was extracted from each rectal swab by mechanical disruption in phenol: chloroform. The DNA was purified using a QIAquick 96 PCR purification kit (Qiagen, MD, USA). Purification was performed per the manufacturer’s instruction with the following minor modifications: (i) sodium acetate (3 M, pH 5.5) was added to Buffer PM to a final concentration of 185 mM to ensure optimal binding of genomic DNA to the silica membrane; (ii) crude DNA was combined with 4 volumes of Buffer PM (rather than 3 volumes); and (iii) DNA was eluted in 100 μL Buffer EB (rather than 80μL). All sequencing was performed at DNA Sequencing Innovation Laboratory at the Center of Genome Sciences and Systems Biology at Washington University in St. Louis. Phased, bi-directional amplification of the v4 region (515-806) of the 16S rRNA gene was employed to generate indexed libraries for Illumina sequencing as described in Faith et al. (2013). Sequencing was performed on an Illumina MiSeq instrument (Illumina, Inc. San Diego, USA), generating 250 bp paired-end reads.

### 16S rRNA gene sequencing and quality control of data

Pairs of 16S rRNA gene sequences were first merged into a single sequence using FLASH v1.2.11 (Magoc and Salzberg, 2011) with a required overlap of at least 100 and less than 250 base pairs in order to provide confident overlap. Sequences with a mean quality score below Q35 were then filtered out using PRINSEQ v0.20.4 (Schmieder and Edwards, 2011). Sequences were oriented in the forward direction and any primer sequences were matched and trimmed off. Mismatch was allowed up to 1. Sequences were subsequently demultiplexed using QIIME v1.9 (Caporaso et al., 2010). Sequences with greater than 97% nucleotide sequence were clustered into operational taxonomic units (**OTU**) using QIIME with the following settings: `max_accepts = 50, max_rejects = 8, percent_subsample = 0.1 and - suppress_step4`. A modified version of GreenGenes (Ley et al., 2006; Schloss and Handelsman, 2006) was used as reference database. Input sequences that had 10% of the reads with no hit to the reference database were then clustered de novo with UCLUST (Schloss and Handelsman, 2006) to generate new reference OTU to which the remaining 90% of reads were assigned. The most abundant sequence in each cluster was used as representative sequence for the OTU. Sparse OTU were then filtered out by requiring a minimum total observation count of 1,200 for an OTU to be retained, the resulting OTU table was rarefied to 10,000 counts per sample and after data processing and quality control 1,755 OTU were retained for further analysis.

### Genotyping

All pigs were genotyped with the PorcineSNP60 v2 BeadChip (Illumina, Inc., San Diego, CA). Quality control procedures were applied by removing the SNPs that had call rate less than 0.90 and minor allele frequency less than 0.05. After quality control the number of SNPs remaining for further analyses was 42,529.

### Phenotypic data

Phenotypic data collection was done as described by Wilson et al. (2016). Meat quality traits (intramuscular fat content (**IMF**), Minolta a* (**MINA**), Minolta b* (**MINB**), minolta L* (**MINL**), ultimate pH (**PH**), subjective color score (**SCOL**), subjective marbling score (**SMARB**), subjective firmness score (**SFIRM**), shearing force (**SSF**)) and carcass composition traits (Belly weight (**BEL**), ham weight (**HAM**), loin weight (**LOIN**), fat depth (**FD**), loin depth (**LD**) and carcass average daily gain (**CADG**)) were used for the current analysis. All the traits were measured as described by Khanal et al. (2019). A summary of traits used in current analysis is reported in Table 1.

**Table 1.**
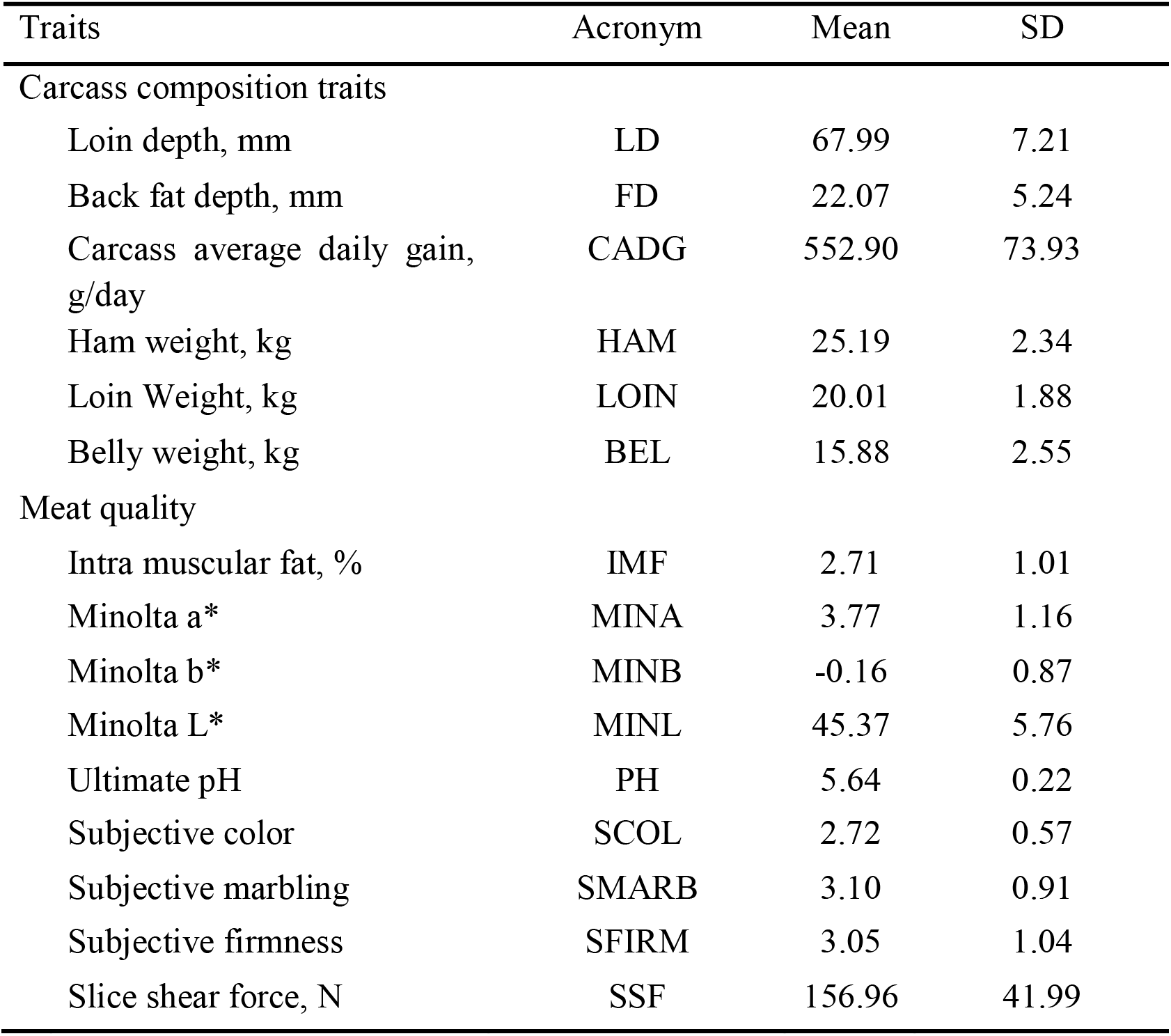
Descriptive statistics of carcass composition and meat quality traits: acronym, means, standard deviation (SD) values.

| Traits | Acronym | Mean | SD |
| --- | --- | --- | --- |
| Carcass composition traits |  |  |  |
| Loin depth, mm | LD | 67.99 | 7.21 |
| Back fat depth, mm | FD | 22.07 | 5.24 |
| Carcass average daily gain, g/day | CADG | 552.90 | 73.93 |
| Ham weight, kg | HAM | 25.19 | 2.34 |
| Loin Weight, kg | LOIN | 20.01 | 1.88 |
| Belly weight, kg | BEL | 15.88 | 2.55 |
| Meat quality |  |  |  |
| Intra muscular fat, % | IMF | 2.71 | 1.01 |
| Minolta a* | MINA | 3.77 | 1.16 |
| Minolta b* | MINB | -0.16 | 0.87 |
| Minolta L* | MINL | 45.37 | 5.76 |
| Ultimate pH | PH | 5.64 | 0.22 |
| Subjective color | SCOL | 2.72 | 0.57 |
| Subjective marbling | SMARB | 3.10 | 0.91 |
| Subjective firmness | SFIRM | 3.05 | 1.04 |
| Slice shear force, N | SSF | 156.96 | 41.99 |

### Statistical analysis

The data were analyzed using ASREML v4.1 (Gilmour et al., 2014). Univariate analyses were conducted to estimate heritabilities, microbiabilities and variance components for each trait. Single trait models were fitted as:

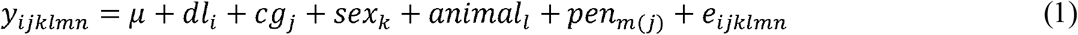

where μ was the overall mean, *dl*_*i*_ was the i^th^ fixed effect of dam line (2 levels), *cg*_*j*_ was the j^th^ fixed effect of the contemporary group (6 levels), *sex*_*k*_ was the k^th^ fixed effect of sex (2 levels), *animail*_*l*_ was the random animal genetic effect, *pen*_*m*(*j*)_ was the random effect of pen nested within contemporary group and *e*_*ijklmn*_ was the random residual. Pen and residuals were assumed normally distributed with mean zero and with variances 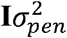 and 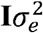, respectively, where **I** was an identity matrix. The random effect of animal was assumed normally distributed with mean 0 and variance 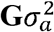 where **G** was a realized genomic relationship matrix obtained according to VanRaden (VanRaden, 2008) as:

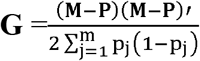

where **M** is a matrix of marker alleles with *m* columns (*m* = total number of markers) and *n* rows (*n* = total number of genotyped individuals), and **P** is a matrix containing the frequency of the second allele (*p*_*j*_), expressed as 2*p*_*j*_. **M**_**ij**_ was −1 if the genotype of individual i for SNP j was homozygous for the first allele, 0 if heterozygous, or 1 if the genotype was homozygous for the second allele. Narrow sense heritabilities were estimated as 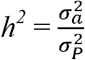, with 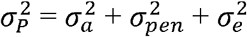.

We added the microbiome information to model (1) in order to estimate the changes in heritability due to the incorporation of microbiome information at each collection stage. Model (2) was then:

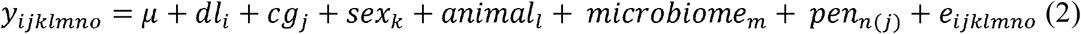

Where *dl*, *cg*, *sex*, *animal*, *pen* and *e* were as previously described and *microniome*_*m*_ was the random effect of the animal microbiome. The effect of the microbiome was assumed normally distributed with mean 0 and variance 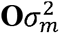 in which **O** was a microbial correlation matrix among individuals and 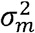 was the microbiome variance. The matrix **O** was created following Camarinha-Silva et al. (2017). Briefly, **O** was obtained as 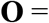 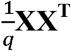, with matrix **X** of dimension of *n* × *q*, where *n* is the number of animals and *q* is the number of OTU. **X** was constructed from **S**, a matrix of equivalent dimensions *n* × *q*. Each element of the **S** matrix, S_*jk*_, was the relative abundance of OTU k in animal j. The elements of X were calculated as:

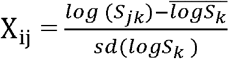

where *S*_*k*_ is the vector of the k^th^ column of **S.** The **O** matrix was created for each stage (Wean, Mid-test and Off-test) separately and fitted in each model separately. The contribution of the microbiome to the overall variance (microbiability) was calculated as: 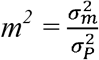 (Difford et al., 2016). The total variance 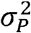 was in this case obtained as 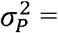 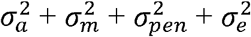.

Bivariate analyses were subsequently conducted to estimate genomic and microbial correlations among traits. Bivariate models were of form:

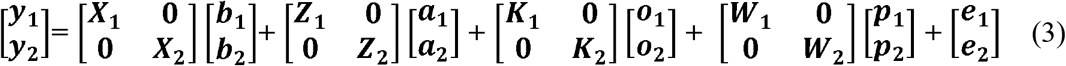

where ***y***_**1**_ and ***y***_**2**_ were the vectors of phenotypic measurements for trait 1 and trait 2 respectively; ***X***_**1**_ and ***X***_**2**_ were the incidence matrices relating the fixed effects to vector ***y***_**1**_ and vector ***y***_**2**_ respectively; ***b***_**1**_ and ***b***_**2**_ were the vector of fixed effect for trait 1 and trait 2 respectively; ***Z***_**1**_ and ***Z***_**2**_ were the incidence matrices relating the phenotypic observations to the vector of random animal effects for trait 1 and trait 2 respectively; ***a***_**1**_ and ***a***_**2**_ were the vectors of random animal effect for trait 1 and trait 2 respectively; ***K***_**1**_ and ***K***_**2**_ were the incidence matrices relating the phenotypic observations to the vector of random microbiome effect for trait 1 and trait 2 respectively; ***o*_1_** and ***o***_**2**_ were the vectors of random microbiome effect for trait 1 and trait 2 respectively; ***W***_**1**_ and ***W***_**2**_ were the incidence matrices relating the phenotypic observations to the vector of random pen effects for trait 1 and trait 2 respectively; ***p***_**1**_ and ***p***_**2**_ were the vector of random pen effect for trait 1 and trait 2 respectively; and ***e***_**1**_ and ***e***_**2**_ were the vectors of random residuals for trait 1 and trait 2 respectively. The fixed effects and random effects were the same as fitted in the univariate analyses.

The additive effects were again assumed normally distributed with means 0 and variance 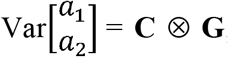; where 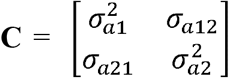. The elements of the covariance matrix **C** were defined as: 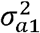 the genetic variance for trait 1, 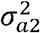, the genetic variance for trait 2, σ_*a*12_ = σ_*a*12_, the additive genetic covariance between trait 1 and trait 2. Similar assumptions were made for the microbiome effect for which the covariance structure was assumed 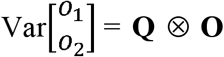; with 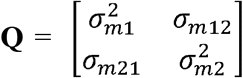. The elements **Q** were: 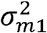, the microbiome variance for trait 1, 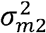, the microbiome variance for trait 2 and *σ*_*m*12_ = *σ*_*m*21_ the microbiome covariance between trait 1 and trait 2. The pen (co)variance structure was 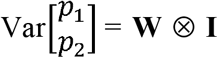; with 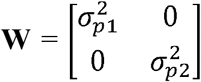 and **I** an identity matrix. The **W** matrix elements were: 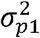, and 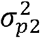 being the pen variance for trait 1 and 2, respectively. Pen effects were in thes case was assumed uncorrelated among traits. The residual variance was given by 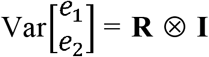; where 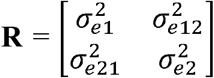 and **I** and was an identity matrix. The components of **R** were defined as: 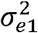 was the residual variance for trait 1, 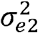 was the residual variance for trait 2, 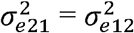 was the residual covariance among the two traits. Preliminary analyses (data not shown), showed how correlations were not estimable for the traits with estimated of microbiome variance of less than 3%. Microbial correlations were therefore estimated among traits for which microbiome explained at least 3% of total phenotypic variance. In all cases microbial correlations were not estimated for weaning since microbiome accounted for less than 3% of total variance for all traits.

### Diversity analysis and its correlations with traits

The diversity analysis performed in this paper was aimed at investigating the distribution of alpha diversities at Wean, Mid-test and Off-test. The R package “vegan” (Oksanen et al., 2019) was used to calculate alpha diversity at each stage. The diversity was measured using Shannon index, and was computed as: 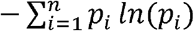 where *p*_*i*_ was the proportional abundance of i^th^ OTU. To estimate the correlation among different traits and alpha diversity at weaning (**alpha_w**), week 15 (**alpha_mid**) and end of test (**alpha_off**), bivariate analyses were conducted using ASREML v.4.1 (Gilmour et al., 2014) by removing the effect of microbiome from model (3) and fitting diversity as the dependent variable.

## Result and discussions

### Data summary, distribution of alpha diversities and variance contributed by each sample

Mean and standard deviation for each meat quality and carcass composition trait are provided in Table 1. There were 9 meat quality and 6 carcass composition traits. The number of individual samples with complete genotypic, phenotypic and microbiome information at each stage was 1,123, which was used for further analyses. The distribution of OTU at weaning, Mid-test and Off-test is given in Figure. 1A. Of a total 1,755 OTU, there were 1,580 OTU in common between weaning, Mid-test and Off-test. There were 1,685 OTU in common between Mid-test and Off-test, while between weaning and Mid-test were 1,626 and between weaning and Off-test were 1590.

**Figure 1.**
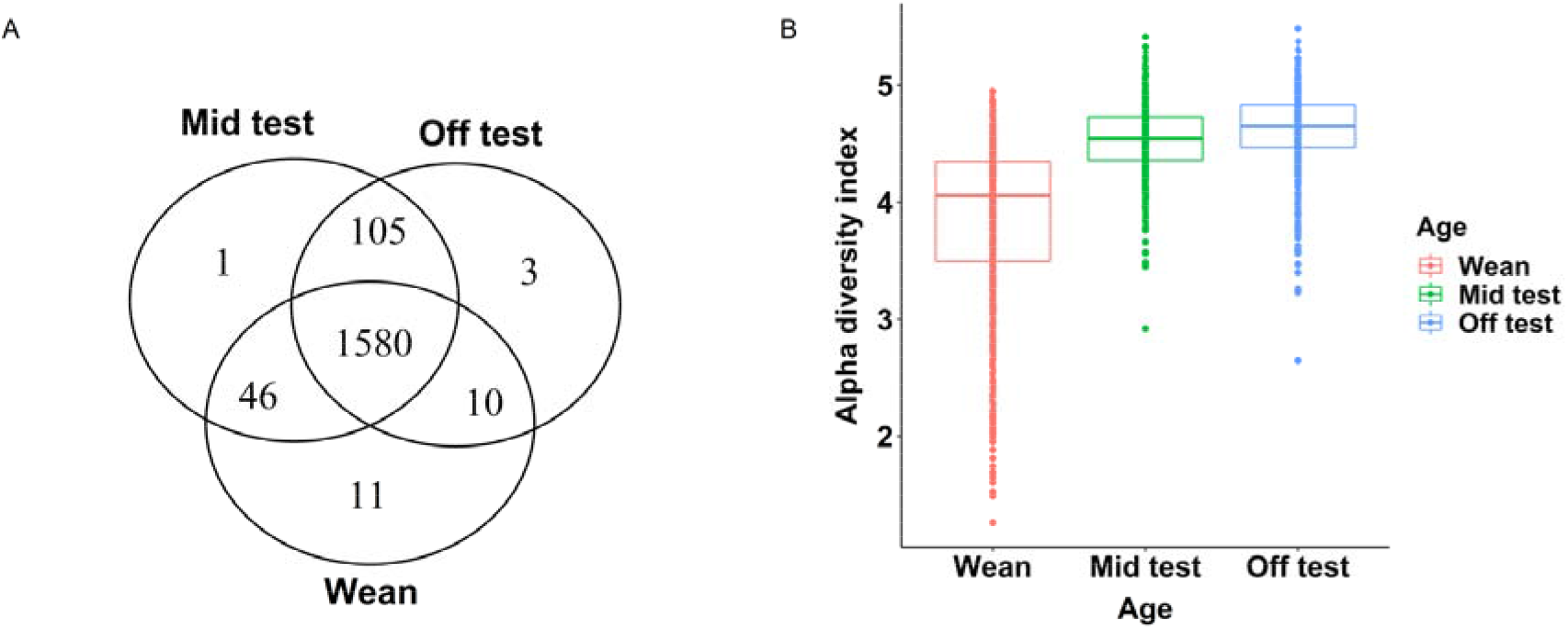
(A) Venn diagram with the numbers of common operational taxonomic units (OTU) among weaning, mid test and off test. (B) Distribution of alpha diversity index among weaning, mid test and off test. X- axis represents the different age group and Y- axis represent the alpha diversity index of each sample for each group.

Alpha diversity is a measure of within-sample diversity. It measures the richness of species and is measured as the number of species in a sample of standard size (Whittaker, 1972). Distribution of alpha diversity among weaning, Mid-test and Off-test is given in Figure 1B. Mean alpha diversity at Off-test, Mid-test and Wean was 4.63 ± 0.01, 4.53 ± 0.01 and 3.85 ± 0.02 respectively. Results from Man-Whitney tests showed that alpha diversity at all stages were different (*P* < 0.001) to each other. This was in accordance with similar studies in pigs and other organisms (Frese et al., 2015; Guevarra et al., 2019); Lu et al., 2018). The increase in alpha diversity with age was similar to what previously found by different authors (Chen et al., 2017; Kim et al., 2011; Looft and Allen, 2012; Thompson et al., 2008). The change in diet in piglet from sow’s milk to complete feed-based diet partially explains the shift in microbial diversity after weaning.

Different researchers (Frese et al., 2015; Konstantinov et al., 2006) reported that change in the diet impact significantly the microbiota composition in the gut. Different types of diet at different stage might explain the difference in alpha diversity at each stage. Piglets are exposed to a large number of stressors during weaning which triggered the physiological change in structure and function of intestine (Guevarra et al., 2019)). This change caused the microbial shift after weaning transition (Kim and Isaacson, 2015) and microbial succession continues until microbiota composition reaches to climax community (Chen et al., 2017) which consists of microbes that are stable in composition. Further higher granularity results on the characterization of the microbial composition in the individuals of the current study can be found in Lu et al. (2018).

### Microbiability estimates

The proportion of variance explained by each random term for meat quality and carcass composition traits is presented in Figure 2 and Figure 3, respectively. The estimates of microbiability and variance components along with their respective standard errors are provided (see additional File 4). The results identified several traits with significant microbiability.

**Figure 2.**
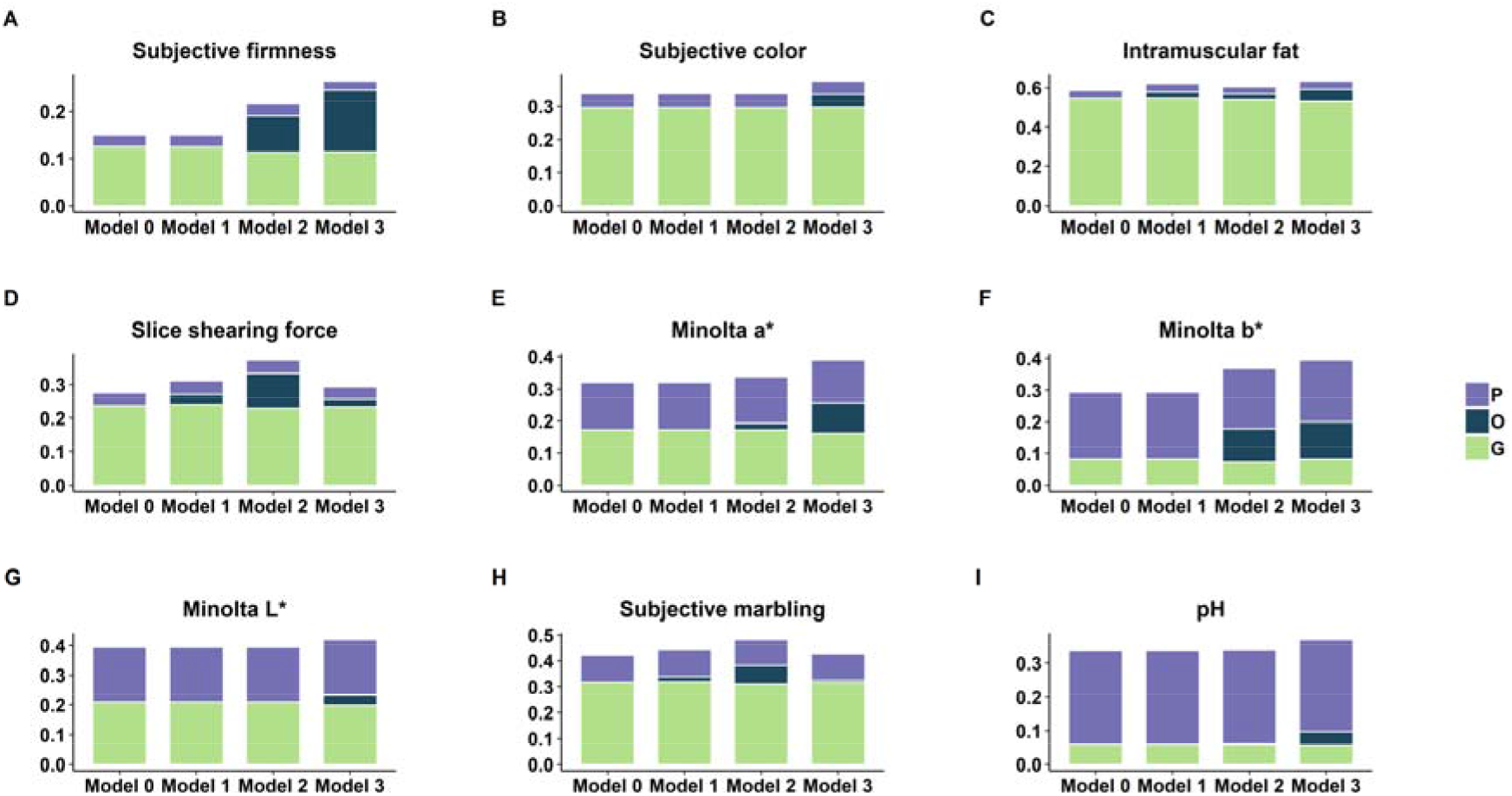
Proportion of variance explained by microbiome relationship matrix (**O**), genomic relationship matrix (**G**) and pen (**P**) for meat quality traits. Model 0 contains **G** matrix and pen effect as random effect, Model 1, Model 2 and Model 3 contains **O** matrix at weaning, Mid-test and Off-test in addition to **G** matrix and pen effect.

**Figure 3.**
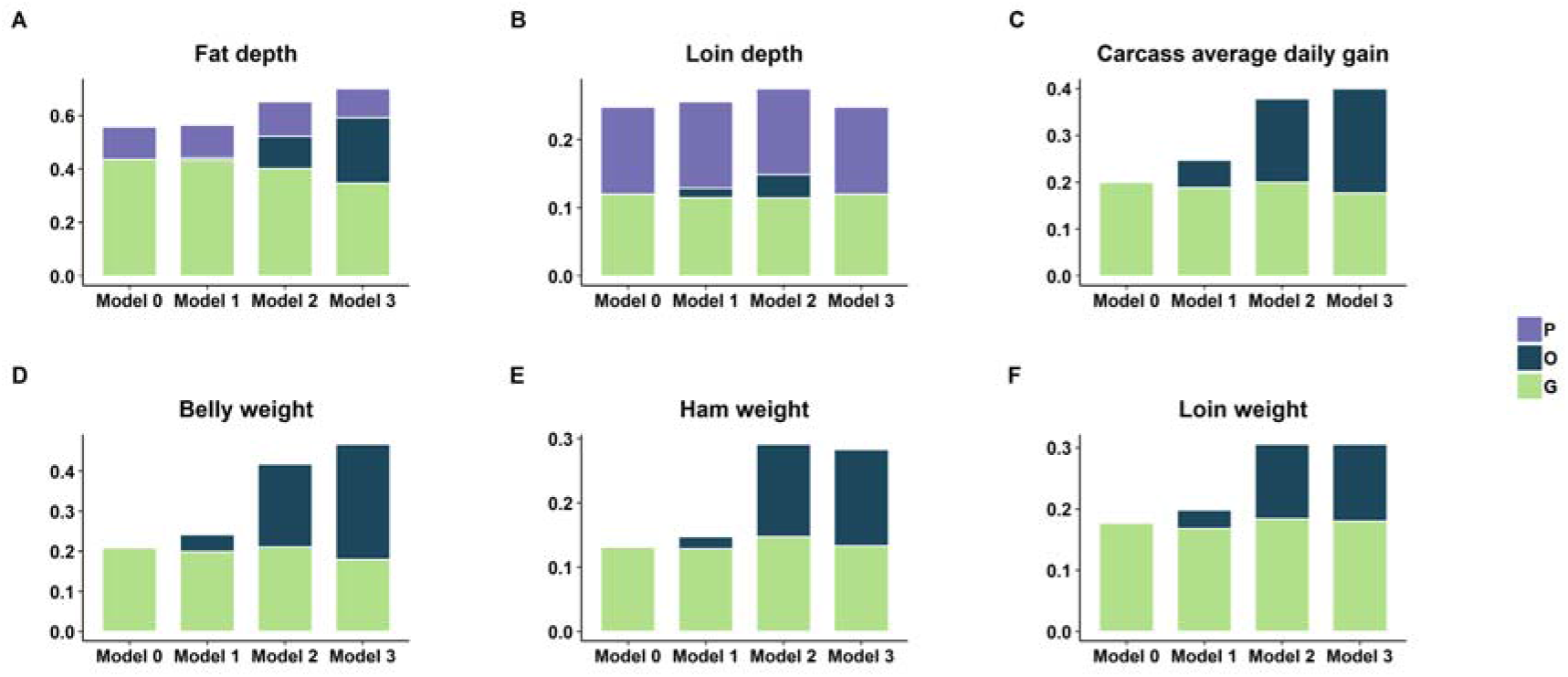
Proportion of variance explained by microbiome relationship matrix (**O**), genomic relationship matrix (**G**) and pen (**P**) for carcass composition traits. Model 0 contains **G** matrix and pen effect as random effect, Model 1, Model 2 and Model 3 contains **O** matrix at weaning, Mid-test and Off-test in addition to G matrix and pen effect.

The microbiability of carcass composition traits were higher than those of meat quality traits. In all cases microbiabilities for both meat quality and carcass composition traits at weaning were negligible and ranged from zero for several traits to a maximum of 0.06 ± 0.03 (estimate ± SE) for CADG. Three of the 9 meat quality traits investigated shown significant microbiability at Mid-test, with estimates of 0.07±0.03 for SMARB, 0.08±0.03 for SFIRM and 0.10±0.04 for MINB. At Off-test, 4 meat quality traits had significant microbiability, with estimates of 0.06±0.02 for IMF, 0.09±0.04 for MINA, 0.11±0.04 for MINB and 0.13±0.04 for SFIRM. For carcass composition traits, we found that 5 out of 6 traits were significantly affected by microbiome at Mid-test and Off-test. The microbiability of carcass composition traits at Mid-test ranged from 0.12 ± 0.04 for LOIN and FD to 0.20 ± 0.04 for BEL. The microbiability of carcass composition traits at Off-test ranged from 0.13 ± 0.05 for LOIN to 0.29 ± 0.05 for BEL. In our study, the microbiome did not contribute significantly to loin depth variability. In most of the cases microbiome at weaning did not contribute to trait variation, however, microbiome at Mid-test and Off-test contributed significantly to trait variation. This might have several causes including the sudden change of microbiome composition shortly after the diet switch occurring at weaning as well as other environmental factors like, stress. To our knowledge this is the first attempt to obtain microbiability estimates for meat quality and carcass composition traits. We did not find any literature to compare the estimates with previous research. Our results suggest that later measures of microbial composition might be more informative for selection purposes, but further research would be needed to clarify this aspect.

Among meat quality traits, microbial variance explained a larger proportion of phenotypic variance than additive genetic for SFIRM and MINB at Off-test (Figure 2). Among carcass composition traits, BEL, HAM, and CADG at Off-test had higher proportion of phenotypic variation explained by microbiome than by additive genetic (Figure 3). These results indicated that a significant proportion of total variance is explained by the microbiome, in some cases larger than the additive genetics and that prediction for these traits could be improved by accounting the effect of variability in gut microbiome composition. The variation in gut microbiome could be fitted as the systematic environmental effect in model.

In the current study we observed a decrease in genomic heritability for most of the carcass composition traits at Off-test when microbiome information was added. The decrease in heritability ranged from 0% for LD to about 10% for FD. At Mid-test, the decrease in heritability ranged from 0% for CADG, BEL, HAM and LOIN to 4% for FD. No change in genomic heritability were observed at weaning. The decrease in heritability for FD was similar to what found by Lu et al. (2018) for similar traits. He et al. (2016) also reported the significant contribution of microbiome in porcine fatness. These results suggested that part of the resemblance among individuals captured by genetic effects in breeding values prediction, might be in fact a resemblance among microbial composition and genetic parameters might not be accurate.

In contrast, for most of the meat quality traits considered, the inclusion of microbial composition did not affect the estimates of genomic heritability, thus suggesting that at least for meat quality traits, gut microbial composition is mostly an environmental factor. The decrease in genomic heritability as we included the microbiome composition in the models was previously observed by Sandoval-Motta et al. (2017) who reported the possibility of overestimation of heritability values with the use of genetic similarities by kinship information. The authors also suggested that inclusion of genetic diversity of individual microbiome will most likely increase the accuracy of heritability of various traits. The heritability and microbiability estimation of daily gain, feed intake and feed conversion ratio in swine by Camarinha-Silva et al. (2017) and methane emission in cattle by Difford et al. (2018) strongly suggested the significant contribution of microbiome in the total variation in the complex phenotypes of livestock. In human, Richards et al. (2018) reported that host genes are affected by the microbiome and are involved in the complex traits. These previous studies agreed with our results. Our results also agreed with Zilber-Rosenberg & Rosenberg, (2008) who reported the concept of “hologenome” of evolution, where the animal or plant along with associated microorganisms are the unit of selection in evolution.

### Correlation of meat quality and carcass composition traits with alpha diversity at different stages

Host genetics plays a major role in shaping the intestinal microbiota of mice and humans (Büsing and Zeyner, 2015; Dąbrowska and Witkiewicz, 2016; Hancox et al., 2015). Chen et al. (2018), Kubasova et al. (2018) and Lu et al. (2018) reported the impact of host genetics on development of gut microbiota in pigs. So, the alpha diversities at weaning, Mid-test and Off-test were considered as separate phenotypic records and genetic correlations were estimated between different alpha diversities and other traits measured. The results are presented in Table 2 suggesting very weak correlations for alpha_w for all traits measured. Weak correlations were estimated between also with alpha_mid with the exception of MINA (−0.45±0.19) where greater alpha diversity seems linked to a paler red color of meat given that MINA is related to the amount of myoglobin in muscle. We obtained weak correlations between alpha_mid and carcass composition traits except for CADG (−0.43±0.19), suggesting that an increase in microbial diversity would decrease average daily gain. This was in contrast with general opinion that the diversity will increase the metabolite production from different microbiota (Kim and Isaacson, 2015; Yan et al., 2017) and increase the weight of host. However, this was in agreement with what found by Lu et al. (2018). Alpha diversity could then be used as a potential indicator trait in CADG selection. In all cases correlations of alpha_off with growth, carcass and meat quality traits were weak (Table 2).

**Table 2.**
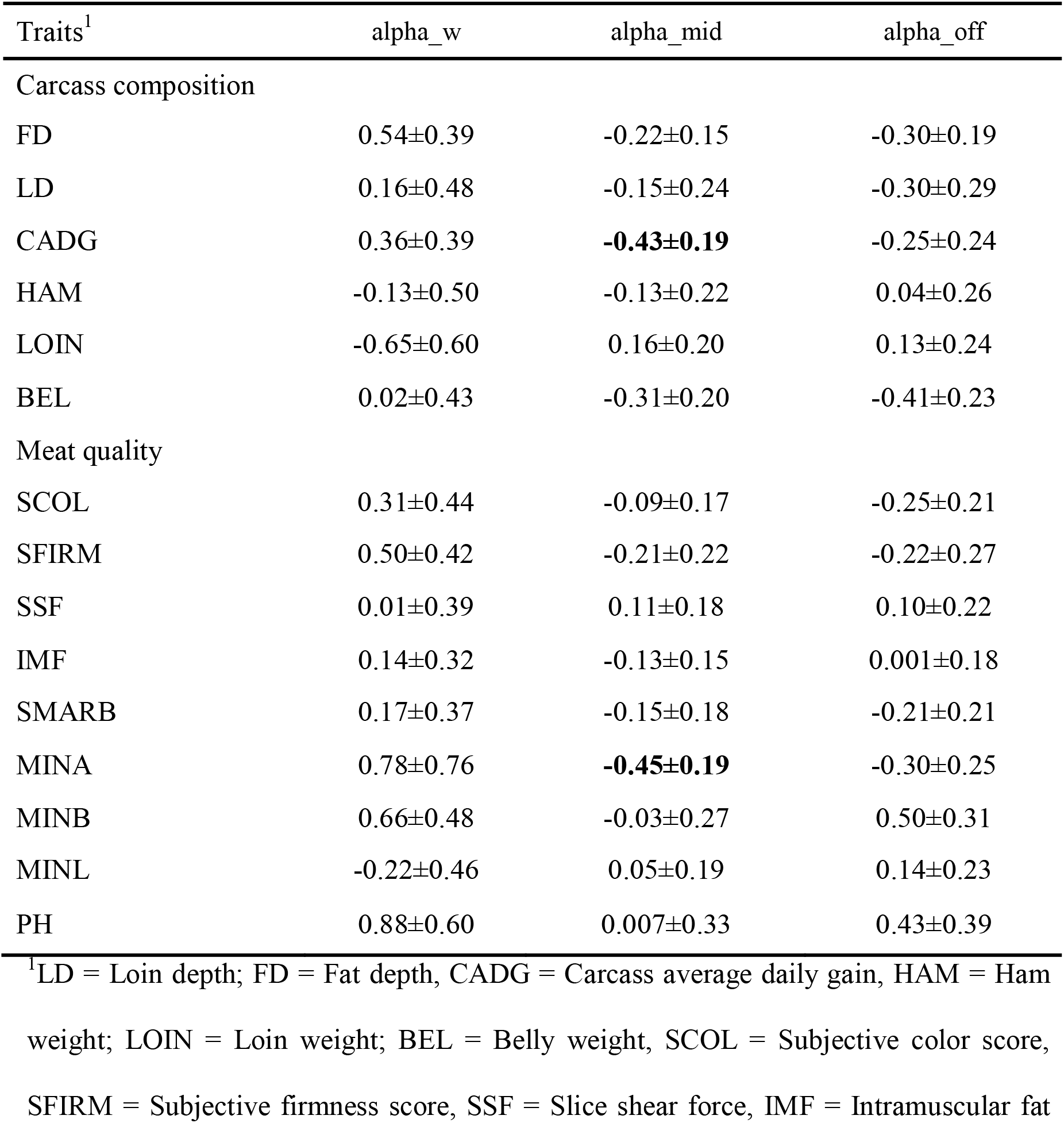
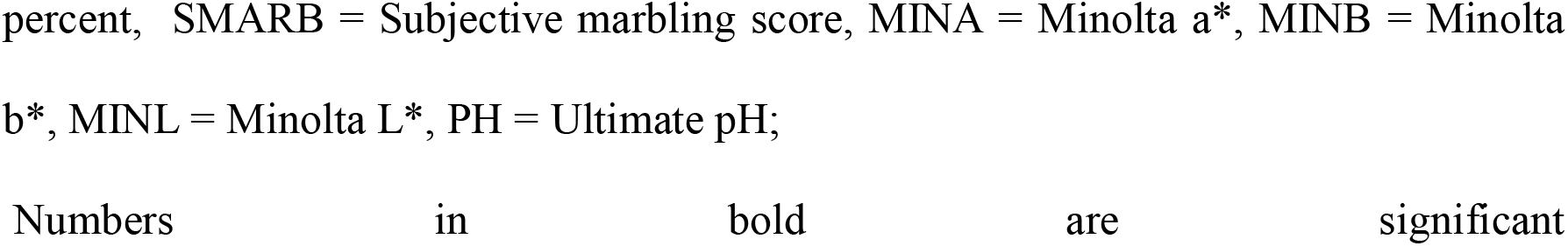
Genetic correlation of carcass composition traits and meat quality traits with alpha diversity at weaning (alpha_w), week 15 (alpha_mid) and end of test (alpha_off).

| Traits<sup>1</sup> | alpha_w | alpha_mid | alpha_off |
| --- | --- | --- | --- |
| Carcass composition |  |  |  |
| FD | 0.54±0.39 | -0.22±0.15 | -0.30±0.19 |
| LD | 0.16±0.48 | -0.15±0.24 | -0.30±0.29 |
| CADG | 0.36±0.39 | **-0.43±0.19** | -0.25±0.24 |
| HAM | -0.13±0.50 | -0.13±0.22 | 0.04±0.26 |
| LOIN | -0.65±0.60 | 0.16±0.20 | 0.13±0.24 |
| BEL | 0.02±0.43 | -0.31±0.20 | -0.41±0.23 |
| Meat quality |  |  |  |
| SCOL | 0.31±0.44 | -0.09±0.17 | -0.25±0.21 |
| SFIRM | 0.50±0.42 | -0.21±0.22 | -0.22±0.27 |
| SSF | 0.01±0.39 | 0.11±0.18 | 0.10±0.22 |
| IMF | 0.14±0.32 | -0.13±0.15 | 0.001±0.18 |
| SMARB | 0.17±0.37 | -0.15±0.18 | -0.21±0.21 |
| MINA | 0.78±0.76 | **-0.45±0.19** | -0.30±0.25 |
| MINB | 0.66±0.48 | -0.03±0.27 | 0.50±0.31 |
| MINL | -0.22±0.46 | 0.05±0.19 | 0.14±0.23 |
| PH | 0.88±0.60 | 0.007±0.33 | 0.43±0.39 |

<sup>1</sup>LD = Loin depth; FD = Fat depth, CADG = Carcass average daily gain, HAM = Ham weight; LOIN = Loin weight; BEL = Belly weight, SCOL = Subjective color score, SFIRM = Subjective firmness score, SSF = Slice shear force, IMF = Intramuscular fat
percent, SMARB = Subjective marbling score, MINA = Minolta a\*, MINB = Minolta b\*, MINL = Minolta L\*, PH = Ultimate pH; Numbers in bold are significant

This study is the first to estimate the genetic and phenotypic correlation between alpha diversity, and carcass and meat quality traits. Our results suggested that diversity at weaning might not be an accurate predictor of growth, carcass and meat quality traits which agreed with Huttenhower et al. (2012). Alpha diversity was reported to be associated with gut health of animal and associated with the normal physiology of host animals (Guevarra et al., 2019)). The major role could include the normal function of gut, enhance immune response and play active role in digestion and utilization of nutrients. Our results presented the varied range of correlation in terms of magnitude and direction at different stages. So, for routine use of the alpha diversity as indicator trait, further investigation of alpha diversity after weaning of piglets is warranted.

### Correlation among traits

In the discussion of correlation, we only focus on microbial correlations. Genomic correlations are only discussed if the genomic correlations changed due to inclusion of microbiome information in the model. The genomic correlations without inclusion of microbiome in the model are presented in additional file 5.

### Correlations among meat quality and carcass composition traits at mid test

Overall there were 3 meat quality traits and 5 carcass composition traits having variance of microbiome composition greater than 3%. Microbial correlations among meat quality traits at Mid-test are presented in Table 3. Most of the microbial correlations were significant. Subjective marbling score was moderately positively correlated (0.46±0.24) with FD. This suggested that shifting of microbiota for high marbled meat would results in higher fat depth. Shear force is the measure of tenderness. In this study, the microbial composition of SSF was highly negatively correlated with SMARB, SFIRM, FD, CADG, LOIN and BEL which ranged from −0.93±0.11 for SSF and SFIRM to −0.50±0.25 for SSF and LOIN. High positive correlations of SFIRM were found with CADG, HAM, LOIN and BEL which ranged from 0.58±0.26 between SFIRM and LOIN to 0.87±0.16 between SFIRM and BEL. We did not find any other estimates to compare with our values of microbial correlation between meat quality and carcass composition traits. There were moderate to high correlations of microbial composition of FD with CADG, HAM, LOIN and BEL which ranged from 0.44±0.21 between FD and LOIN to 0.74±0.11 between FD and BEL. High positive correlations were found between CADG and HAM, LOIN and BEL. Belly weight was highly positively correlated with HAM (0.96±0.03) and LOIN (0.94±0.06).

**Table 3.** Estimates of microbial correlation (above diagonal) and genomic correlation (below diagonal) at Mid-test among meat quality and carcass composition traits.

|  | <sup>1</sup>SMARB | SFIRM | SSF | FD | CADG | HAM | LOIN | BEL |
| --- | --- | --- | --- | --- | --- | --- | --- | --- |
| **SMARB** |  | 0.39±0.33 | **-0.72±0.28** | **0.46±0.24** | -0.21±0.28 | -0.27±0.29 | -0.34±0.32 | -0.02±0.26 |
| **SFIRM** | **0.42±0.18** |  | **-0.93±0.11** | NC<sup>2</sup> | **0.86±0.17** | **0.62±0.24** | **0.58±0.26** | **0.87±0.16** |
| **SSF** | 0.08±0.16 | -0.23±0.21 |  | **-0.70±0.21** | **-0.68±0.22** | -0.45±0.25 | **-0.50±0.25** | **-0.55±0.24** |
| **FD** | **0.22±0.11** | NC | **-0.44±0.13** |  | **0.68±0.15** | **0.50±0.19** | **0.44±0.21** | **0.74±0.11** |
| **CADG** | 0.02±0.17 | 0.03±0.23 | 0.19±0.18 | 0.21±0.15 |  | **0.98±0.02** | **0.95±0.03** | **0.98±0.01** |
| **HAM** | -0.13±0.18 | 0.11±0.24 | 0.27±0.20 | 0.01±0.15 | **0.67±0.11** |  | NE<sup>3</sup> | **0.96±0.03** |
| **LOIN** | -0.09±0.17 | 0.10±0.23 | 0.11±0.18 | -0.14±0.15 | **0.69±0.09** | **0.53±0.11** |  | **0.94±0.06** |
| **BEL** | 0.31±0.17 | 0.35±0.23 | 0.18±0.15 | **0.57±0.11** | **0.79±0.06** | **0.42±0.17** | **0.42±0.15** |  |

<sup>1</sup>SMARB = Subjective marbling score, SFIRM = Subjective firmness score, SSF = Slice shear force, FD = Fat depth,
CADG = Carcass average daily gain, HAM = Ham weight; LOIN = Loin weight; BEL = Belly weight;
<sup>2</sup>Not Converged; <sup>3</sup>Not estimable; Numbers in bold are significant

### Correlation between meat quality traits and carcass composition traits at the end of test

There were 6 meat quality traits and 5 carcass composition for which variance of microbiome composition was greater than 3%. The microbial and genomic correlations among meat quality traits at Off-test are presented in Table 4. pH had high positive microbial correlation (0.90±0.25) with SCOL and SFIRM (0.73±0.35). This is in partial agreement with results from Ratzke & Gore, (2018) that reported how there are specific bacteria which build lactic acid in the muscle resulting in the anaerobic breakdown of glucose and glycogen, which eventually loosens the myofibril, thus scattering more light making the muscle pale (Walters, 1975). Furthermore, increasing pH causes swelling of myofibrils (Huff-Lonergan and Lonergan, 2005) which ultimately makes the muscle firmer. High positive microbial correlation was found between IMF and SFIRM (0.91±0.17), MINA (0.55±0.28) and MINB (0.75±0.27). This agrees with Fang, Xiong, Su, Huang, & Chen. (2017) who reported that gut bacteria involving in energy metabolism and intramuscular fat content in pig also regulate the muscle composition and muscle fibers. Higher microbial correlation of IMF with minolta color measurements and SFIRM indicated that microbial composition increasing IMF would make the muscle pale and firmer. High microbial correlation of MINA and MINB (0.78±0.16) suggests that microbiota responsible for redness of meat also contribute to the yellowness in the meat. This agreed with Kim et al. (2010) who reported the positive correlation of yellowness and redness in the muscle of pig.

**Table 4.** Estimates of microbial correlation (above diagonal) and genomic correlation (below diagonal) at end of test among meat quality traits.

|  | <sup>1</sup>SCOL | IMF | SFIRM | MINA | MINB | PH |
| --- | --- | --- | --- | --- | --- | --- |
| **SCOL** |  | -0.28±0.57 | 0.07±0.31 | 0.29±0.44 | -0.26±0.39 | **0.90±0.25** |
| **IMF** | -0.22±0.13 |  | **0.91±0.17** | **0.55±0.28** | **0.75±0.27** | 0.10±0.47 |
| **SFIRM** | 0.18±0.19 | 0.29±0.17 |  | 0.26±0.27 | 0.12±0.26 | **0.73±0.35** |
| **MINA** | **0.45±0.16** | **0.29±0.14** | -0.53±0.28 |  | **0.78±0.16** | 0.33±0.36 |
| **MINB** | **-0.94±0.22** | **0.78±0.16** | -0.03±0.32 | -0.10±0.27 |  | 0.38±0.38 |
| **PH** | 0.13±0.50 | -0.18±0.25 | 0.44±0.36 | -0.04±0.33 | -0.47±0.42 |  |

<sup>1</sup>SCOL = Subjective color score, SFIRM = Subjective firmness score, IMF = Intramuscular fat percent, MINA = Minolta a\*, MINB = Minolta b\*, PH = Ultimate pH; Numbers in bold are significant.

The microbial and genomic correlations among carcass composition traits at Off-test are presented in Table 5. The microbial composition of carcass composition traits were highly and positively correlated to each other ranging from 0.55±0.17 between FD and LOIN to 0.97±0.02 between CADG and HAM. McCormack et al. (2018) reported a positive correlation between gut microbiota and feed efficiency in swine. Gut microbiota has considerable effect on feed intake, final body weight (Kubasova et al., 2018) and growth traits (Ramayo-Caldas et al., 2016). All these studies suggested that microbial composition has considerable effects on many carcass composition traits, with positive correlations between them. This high correlation indicated that all the traits could be simultaneously improved through the same microbial composition.

**Table 5.** Estimates of microbial correlation (above diagonal) and genomic correlation (below diagonal) at end of test among carcass composition traits.

|  | <sup>1</sup>FD | CADG | HAM | LOIN | BEL |
| --- | --- | --- | --- | --- | --- |
| **FD** |  | **0.71±0.11** | **0.59±0.16** | **0.55±0.17** | **0.94±0.05** |
| **CADG** | 0.14±0.15 |  | **0.97±0.02** | **0.91±0.05** | **0.94±0.03** |
| **HAM** | -0.10±0.17 | **0.63±0.13** |  | <sup>2</sup>NE | **0.87±0.06** |
| **LOIN** | -0.13±0.15 | **0.67±0.10** | **0.54±0.19** |  | **0.82±0.08** |
| **BEL** | **0.49±0.13** | **0.78±0.07** | 0.34±0.19 | **0.40±0.16** |  |

<sup>1</sup>FD = Fat depth, CADG = Carcass average daily gain, HAM = Ham weight; LOIN = Loin weight; BEL = Belly weight; <sup>2</sup>Non estimable; Numbers in bold are significant.

The microbial correlations for meat quality traits and carcass composition traits at Off-test are presented in Table 6. Intramuscular fat was highly correlated with FD (0.90±0.14) and BEL (0.73±0.18). Firmness score was positively correlated with BEL (0.50±0.18). Moderate positive correlation was found between MINA and BEL (0.41±0.21) and high positive correlation was found between MINA and FD (0.53±0.18), and MINA and CADG (0.66±0.17). Minolta b* has moderate positive correlation with FD (0.43±0.19) and high positive correlation with CADG (0.58±0.18): suggesting that increase in microbiota for lean meat and high daily gain of carcass would make the meat more yellowish.

**Table 6.**
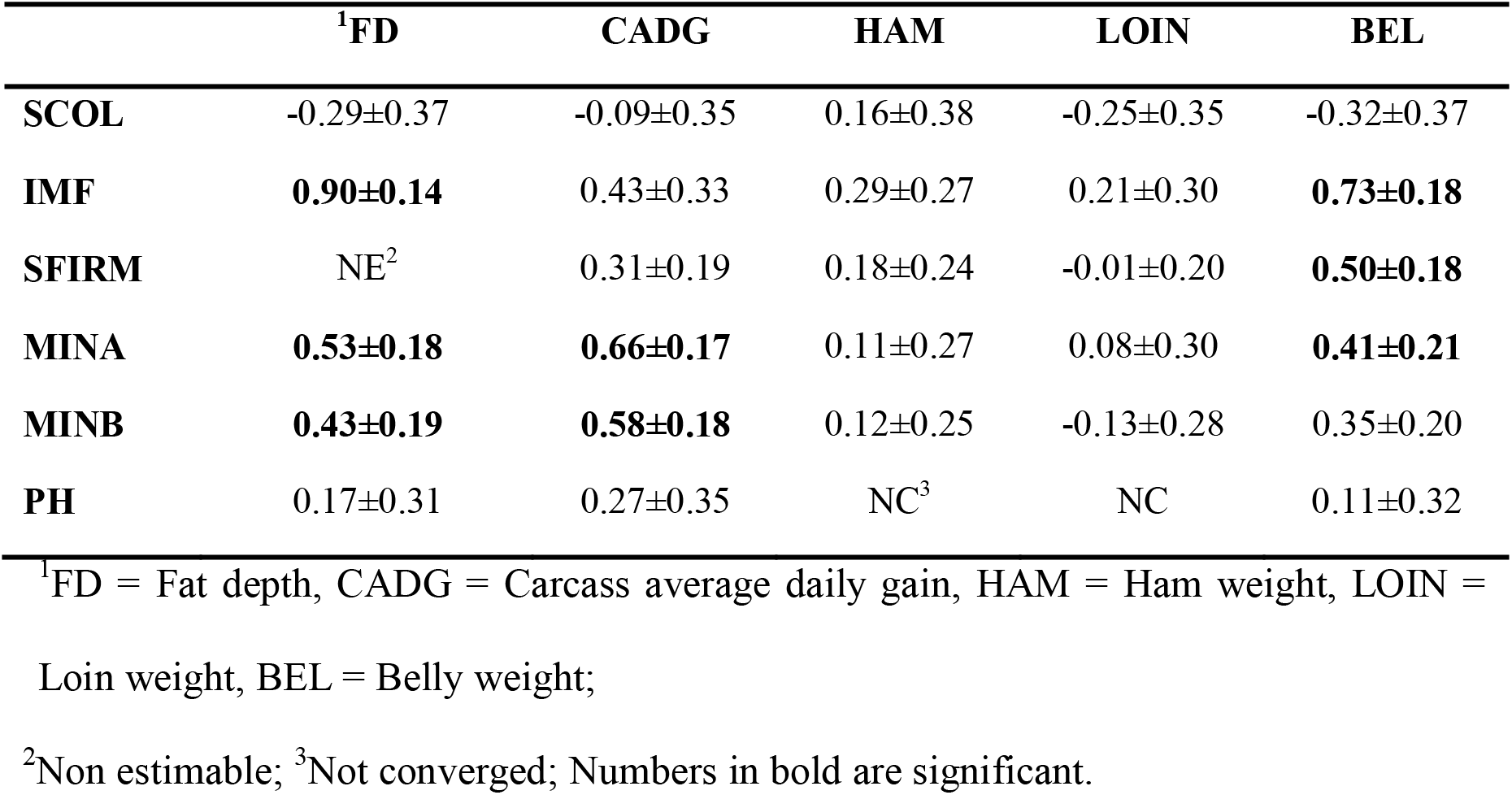
Estimates of microbial correlation between meat quality traits and carcass composition traits at Off test.

|  | <sup>1</sup>FD | CADG | HAM | LOIN | BEL |
| --- | --- | --- | --- | --- | --- |
| **SCOL** | -0.29±0.37 | -0.09±0.35 | 0.16±0.38 | -0.25±0.35 | -0.32±0.37 |
| **IMF** | **0.90±0.14** | 0.43±0.33 | 0.29±0.27 | 0.21±0.30 | **0.73±0.18** |
| **SFIRM** | NE<sup>2</sup> | 0.31±0.19 | 0.18±0.24 | -0.01±0.20 | **0.50±0.18** |
| **MINA** | **0.53±0.18** | **0.66±0.17** | 0.11±0.27 | 0.08±0.30 | **0.41±0.21** |
| **MINB** | **0.43±0.19** | **0.58±0.18** | 0.12±0.25 | -0.13±0.28 | 0.35±0.20 |
| **PH** | 0.17±0.31 | 0.27±0.35 | NC<sup>3</sup> | NC | 0.11±0.32 |

<sup>1</sup>FD = Fat depth, CADG = Carcass average daily gain, HAM = Ham weight, LOIN = Loin weight, BEL = Belly weight;
<sup>2</sup>Non estimable; <sup>3</sup>Not converged; Numbers in bold are significant.

### Change in genomic correlation with the inclusion of microbiome information

In this study, we observed a decrease in genomic correlation for the majority of the carcass composition traits when microbiome information was included in the model. At Mid-test, the decrease in genomic correlation ranged from 0% for majority of meat quality traits to 18% for BEL and LOIN. The genomic correlation of BEL with FD and HAM decreased by 5% and 16%, respectively. The genomic correlation of FD with SMARB and SSF decreased by 7% and 4%, respectively.

At Off-test, the genomic correlation between PH and SCOL (0.91±0.29), SFIRM and IMF (0.36±0.15), FD and CADG (0.27±0.13), and BEL and HAM (0.58±0.19) became non-significant with the inclusion of microbiome. Among carcass traits, the decrease in genomic correlation ranged from 1% between BEL and CADG to 30% between BEL and LOIN. The genomic correlation of BEL with FD, CADG with HAM, CADG with LOIN, FD with IMF, FD with MINB, BEL with IMF, and BEL with SFIRM decreased by 13%, 4%, 2%, 9%, 6%, 13% and 8%, respectively. Among meat quality and carcass traits, the decrease in genomic correlations ranged from 1% for FD and SFIRM to 9% for BEL and IMF. We observed the decrease in genomic correlations with the inclusion of microbiome, particularly of any other traits with fat related traits e.g. (BEL, FD, IMF). This could be due to the greater influence of gut microbiome on fat deposition. Furthermore, we observed that there was a decrease in genomic correlation for those traits which had higher microbial correlation. High microbial correlations among different traits suggested that genomic correlations among traits are partially contributed by the correlations among the gut microbiota composition. The covariance among microbiome for different traits might have contributed to the genetic covariance and hence the genomic correlation. We observed that the decrease in the genomic correlation was higher at Off-test than at Mid-test. This was due to high variability accounted by microbiome composition at Off-test in comparison to Mid-test.

This is the first study to evaluate the variance accounted by microbiome and estimate the microbial correlations for meat quality and carcass traits in swine. So, we have explored the model sequentially, first with inclusion of genomic information and then addition of microbiome information at different stages. Variance component estimates of different random effects with inclusion of interaction of genotype-by-microbiome in the model is recommended for future studies.

## Conclusions

This study was conducted on crossbred pigs to investigate the impact of intestinal microbiota through different stages (weaning, Mid-test and Off-test) of production. To our knowledge this study is the first attempt to investigate the impact of microbiome on the meat quality and carcass composition traits at a large scale in swine. The contribution of microbiome to all traits was significant although it varied over time with an increase from weaning to Off-test for most of the traits. Adding microbiome information did not affect the estimates of genomic heritability of meat quality traits but changed the estimate of carcass composition traits suggesting that portion of genomic variance was contributed by gut microbiome. Alpha diversity at Mid-test was strongly correlated with carcass average daily gain and minolta a* color score. A better understanding of microbial composition could aid the improvement of complex traits, particularly the carcass composition traits in swine by inclusion of microbiome information in the genetic evaluation process. High microbial correlations were found among different traits, particularly with traits related to fat deposition. Adding microbiome information decreased the genomic correlation for those traits which had higher microbial correlation suggesting that portion of genomic correlation was due to genetic covariance among microbiome composition affecting those traits. Based on the results we can conclude that microbial composition could be altered to improve a given trait. To obtain optimum microbial composition, manipulation of gut microbiota could be done using specific bacterial composition as probiotics or increasing the relative abundance through prebiotics, feed additives supplements and fecal microbiota transplantation could also be done. The estimated parameters provide a reference value for further research on gut microbial contribution to complex phenotypes in pigs. These results may lead to establish a newer approach of genetic evaluation process through the addition of gut microbial information.

## Acknowledgements

We would like to thank Jessica Hoisington-Lopez from the DNA sequencing Innovation Lab at the Center for Genome Sciences and Systems Biology at Washington University in St. Loius for her sequencing expertise and Nicholas S. Grohmann for phenotype and sample collection.

## Funding

This study is a part of the project “Re-defining growth efficiency accounting for the interaction between host genome and commensal gut bacteria” funded by The National Pork Board Association and part of the project “From Host to Guest and back” funded by The Maschhoffs, LLC and North Carolina State University.

## Author Contributions

PK designed and carried all analyses, as well as interpreted the results and drafted the manuscript. CS and JF were involved in designing the experiment and helped in interpretation of the results. CM and FT were involved in designing the experiment and providing the consultation for the analyses. All co-authors provided comments for manuscript. All authors have read and approved the final manuscript.

## Competing interests

The authors declare that they have no competing interests.

## Contribution to the field statements

This manuscript describes about the impact of gut microbiome composition at different stages of production on meat quality and carcass composition traits. Until recently, selection of different traits in pigs has been done with the use of pedigree and genomic information, yet the advantage of incorporating microbial information in the genetic evaluation processes has not been assessed. So, this study evaluates the variance accounted by microbial composition and its effect on heritability and genomic correlation. Adding microbiome information did not affect the estimates of genomic heritability of meat quality traits but affected the estimates of carcass composition traits. We found high microbial correlations among several traits which suggested that genomic correlation was partially contributed by genetic similarity of microbiome composition. Since this is one of the earlier studies in determining the effect of microbiome composition in heritability and genomic correlation of meat quality and carcass traits in swine, we believe that these parameters will provide a reference value to for the researchers in future who wants to conduct research on effect of gut microbiome in complex phenotypes of swine.

