## Additional file 1 for "Microbiability of meat quality and carcass composition traits in swine"

Table S1. Diet formulae and their nutritional values

| | Nursery 3 | Nursery 4 | GF-1 | GF-2 | | GF-3 | | GF-4 | | GF-5 | | GF-6 | | GF-7 |
|---|---|---|---|---|---|---|---|---|---|---|---|---|---|---|
| Ingredient | Barrow/Gilt | Barrow/Gilt | Barrow/Gilt | Barrow | Gilt | Barrow | Gilt | Barrow | Gilt | Barrow | Gilt | Barrow | Gilt | Barrow/Gilt |
| Corn | 660.60 | 861.88 | 800.62 | 1020.46 | 1013.36 | 1236.71 | 1204.82 | 1382.01 | 1335.94 | 1481.33 | 1435.26 | 1530.99 | 1499.03 | 1534.50 |
| Corn germ meal | 48.01 | 341.84 | 564.17 | 464.72 | 467.93 | 366.89 | 381.32 | 301.16 | 322.00 | 256.23 | 277.07 | 233.76 | 248.22 | 232.18 |
| Soybean meal | 326.70 | 594.00 | 490.79 | 394.42 | 397.53 | 299.63 | 313.60 | 235.93 | 256.12 | 192.39 | 212.59 | 170.62 | 184.63 | 169.09 |
| Fat - yellow grease (post-pellet) | | | 64.46 | 47.08 | 47.64 | 29.99 | 32.51 | 18.50 | 22.14 | 10.65 | 14.29 | 6.72 | 9.25 | 6.45 |
| Limestone | | 30.20 | 27.99 | 25.81 | 25.88 | 23.66 | 23.97 | 22.21 | 22.67 | 21.23 | 21.69 | 20.73 | 21.05 | 20.70 |
| Pelleting aid | | | 10.00 | 10.00 | 10.00 | 10.00 | 10.00 | 10.00 | 10.00 | 10.00 | 10.00 | 10.00 | 10.00 | 10.00 |
| L-Lysine HCl (98%) | | 8.91 | 9.65 | 8.17 | 8.22 | 6.71 | 6.93 | 5.74 | 6.05 | 5.07 | 5.38 | 4.73 | 4.95 | 4.71 |
| Salt | | 11.17 | 9.14 | 9.13 | 9.13 | 9.12 | 9.13 | 9.12 | 9.12 | 9.11 | 9.12 | 9.11 | 9.11 | 9.11 |
| Fat - yellow grease | 41.55 | 13.86 | 7.00 | 7.00 | 7.00 | 7.00 | 7.00 | 7.00 | 7.00 | 7.00 | 7.00 | 7.00 | 7.00 | 7.00 |
| Monocalcium phosphate (21%) | | 18.13 | 5.51 | 4.52 | 4.55 | 3.54 | 3.69 | 2.89 | 3.09 | 2.44 | 2.65 | 2.21 | 2.36 | 2.20 |
| HMTBa | | 2.26 | 4.50 | 3.29 | 3.33 | 2.09 | 2.27 | 1.29 | 1.55 | 0.74 | 1.00 | 0.47 | 0.65 | 0.45 |
| L-Threonine (98%) | | 2.07 | 2.35 | 1.78 | 1.80 | 1.23 | 1.31 | 0.85 | 0.97 | 0.60 | 0.72 | 0.47 | 0.55 | 0.46 |
| Trace mineral premix | | 1.98 | 2.00 | 1.87 | 1.87 | 1.73 | 1.75 | 1.64 | 1.67 | 1.58 | 1.61 | 1.55 | 1.57 | 1.55 |
| Phytase 2500 | | 1.90 | 0.80 | 0.76 | 0.76 | 0.73 | 0.73 | 0.70 | 0.71 | 0.68 | 0.69 | 0.67 | 0.68 | 0.67 |
| Vitamin premix | | 0.99 | 0.60 | 0.57 | 0.57 | 0.55 | 0.55 | 0.53 | 0.53 | 0.52 | 0.52 | 0.51 | 0.51 | 0.51 |
| Copper chloride (58%) | | 0.68 | 0.43 | 0.43 | 0.43 | 0.43 | 0.43 | 0.43 | 0.43 | 0.43 | 0.43 | 0.43 | 0.43 | 0.43 |
| Nursery basemix | 791.99 | | | | | | | | | | | | | |
| DDGS | 111.15 | 76.52 | | | | | | | | | | | | |
| Mecadox 2.5 (g/lb) | 20 | 20 | | | | | | | | | | | | |
| Zinc oxide (72%) | | 6.93 | | | | | | | | | | | | |
| Organic acidifier | | 5.94 | | | | | | | | | | | | |
| Carbohydrase | | 0.74 | | | | | | | | | | | | |
| Total: | 2000 | 2000 | 2000 | 2000 | 2000 | 2000 | 2000 | 2000 | 2000 | 2000 | 2000 | 2000 | 2000 | 2000 |

| Nutrient | Units | Nursery 3 | Nursery 4 | GF-1 | GF-2 Barrow | GF-2 Gilt | GF-3 Barrow | GF-3 Gilt | GF-4 Barrow | GF-4 Gilt | GF-5 Barrow | GF-5 Gilt | GF-6 Barrow | GF-6 Gilt | GF-7 |
|---|---|---|---|---|---|---|---|---|---|---|---|---|---|---|---|
| Metabolizable energy | Kcal/lb | 1500 | 1519.371 | 1460.004 | 1460.18 | 1460.17 | 1460.35 | 1460.33 | 1460.47 | 1460.43 | 1460.55 | 1460.51 | 1460.59 | 1460.56 | 1460.59 |
| Crude protein | % | 20.103 | 22.561 | 21.281 | 18.59 | 18.68 | 15.95 | 16.34 | 14.17 | 14.74 | 12.96 | 13.52 | 12.35 | 12.74 | 12.31 |
| Cystine, Dig | % | 0.293 | 0.272 | 0.244 | 0.22 | 0.22 | 0.20 | 0.20 | 0.19 | 0.19 | 0.18 | 0.18 | 0.17 | 0.18 | 0.17 |
| Isoleucine, Dig | % | 0.675 | 0.786 | 0.715 | 0.62 | 0.62 | 0.52 | 0.53 | 0.45 | 0.47 | 0.41 | 0.43 | 0.39 | 0.40 | 0.38 |
| Lysine, Total | % | 1.405 | 1.521 | 1.454 | 1.23 | 1.24 | 1.01 | 1.05 | 0.87 | 0.91 | 0.77 | 0.81 | 0.72 | 0.75 | 0.71 |
| Lysine, Dig | % | 1.25 | 1.34 | 1.27 | 1.07 | 1.08 | 0.87 | 0.90 | 0.74 | 0.78 | 0.65 | 0.69 | 0.61 | 0.64 | 0.60 |
| Leucine, Dig | % | 1.469 | 1.586 | 1.439 | 1.31 | 1.31 | 1.18 | 1.20 | 1.09 | 1.12 | 1.03 | 1.06 | 1.00 | 1.02 | 1.00 |
| Met + Cys, Dig | % | 0.707 | 0.765 | 0.725 | 0.62 | 0.62 | 0.51 | 0.53 | 0.44 | 0.47 | 0.40 | 0.42 | 0.37 | 0.39 | 0.37 |
| Threonine, Dig | % | 0.76 | 0.804 | 0.762 | 0.65 | 0.65 | 0.54 | 0.55 | 0.46 | 0.48 | 0.41 | 0.43 | 0.38 | 0.40 | 0.38 |
| Tryptophan, Dig | % | 0.223 | 0.232 | 0.216 | 0.18 | 0.18 | 0.15 | 0.16 | 0.13 | 0.14 | 0.12 | 0.12 | 0.11 | 0.11 | 0.11 |
| Valine, Dig | % | 0.826 | 0.871 | 0.826 | 0.72 | 0.72 | 0.62 | 0.63 | 0.55 | 0.57 | 0.50 | 0.52 | 0.47 | 0.49 | 0.47 |
| Phosphorus | % | 0.728 | 0.689 | 0.557 | 0.50 | 0.50 | 0.44 | 0.45 | 0.41 | 0.42 | 0.38 | 0.39 | 0.37 | 0.38 | 0.37 |
| P, Available | % | 0.569 | 0.4 | 0.3 | 0.27 | 0.27 | 0.24 | 0.24 | 0.21 | 0.22 | 0.20 | 0.21 | 0.19 | 0.20 | 0.19 |
| Calcium | % | 0.809 | 0.896 | 0.7 | 0.63 | 0.63 | 0.57 | 0.58 | 0.52 | 0.54 | 0.49 | 0.51 | 0.48 | 0.49 | 0.48 |
| Moisture | % | 13.735 | 13.035 | 13.172 | 13.60 | 13.58 | 14.01 | 13.95 | 14.29 | 14.20 | 14.48 | 14.39 | 14.58 | 14.52 | 14.58 |
| Crude fat | % | 5.476 | 3.03 | 5.458 | 4.82 | 4.84 | 4.18 | 4.28 | 3.76 | 3.89 | 3.47 | 3.60 | 3.32 | 3.42 | 3.31 |
| Crude fiber | % | 2.117 | 3.18 | 3.486 | 3.17 | 3.18 | 2.85 | 2.90 | 2.64 | 2.71 | 2.50 | 2.57 | 2.43 | 2.47 | 2.42 |
| ADF | % | 2.92 | 5.01 | 5.367 | 4.86 | 4.88 | 4.37 | 4.44 | 4.03 | 4.14 | 3.81 | 3.91 | 3.69 | 3.76 | 3.68 |
| NDF | % | 6.507 | 12.75 | 15 | 13.58 | 13.63 | 12.19 | 12.40 | 11.26 | 11.55 | 10.62 | 10.91 | 10.30 | 10.50 | 10.28 |
