## Additional file 2 for "Microbiability of meat quality and carcass composition traits in swine"

Table S2. Vaccinations

| Condition | Timing |
|---|---|
| Mycoplasma Hyopneumoniae | Processing (~4 days old) |
| Porcine Circovirus Type 2 (PCV2) | Weaning |
| Porcine Respiratory Syndrome (PRRS) | 10-14 days post-weaning |
| Porcine Circovirus | 14-17 days post PRRS |
| Mycoplasma Hyopneumoniae | vaccination |
| Ileitis | ~6 weeks post-weaning |
| Erysipelas | |

Table S3. Injectable medications

| Condition | Product | Timing |
|---|---|---|
| Respiratory, Diarrhea, Lameness | Excede | Weaning to 8 weeks post-weaning |
| Respiratory | Biomycin 200 | 8-14 weeks post-weaning |
| Respiratory | Lincocin 300 | 14 weeks post-weaning to end of study |
| Diarrhea, Lameness | Lincocin 300 | 8 weeks post-weaning to end of study |
| Lameness | Dexamethasone | Weaning to 14 weeks post-weaning |

Table S4. Water medications

| Condition | Product | Timing |
|---|---|---|
| Diarrhea | Neomycin | Weaning |
| Respiratory, Diarrhea | Oxytetracycline (OTC)<br>Denagard | As Needed |
| Respiratory, Diarrhea, Lameness | Linco Soluble | As Needed |
