## Additional file 3 for "Microbiability of meat quality and carcass composition traits in swine"

Table S5. Distribution of samples across families, sex, and time points

|  | | Female |  |  |  | Male |  | Total |
| --- | --- | --- | --- | --- | --- | --- | --- | --- |
| Family | Weaning | Mid-test | Off-test |  | Weaning | Mid-test | Off-test |  |
| 1 | 22 | 23 | 22 |  | 20 | 20 | 20 | 127 |
| 2 | 19 | 25 | 24 |  | 23 | 23 | 21 | 135 |
| 3 | 20 | 23 | 22 |  | 20 | 23 | 23 | 131 |
| 4 | 23 | 24 | 23 |  | 21 | 23 | 23 | 137 |
| 5 | 21 | 18 | 21 |  | 15 | 15 | 15 | 105 |
| 6 | 21 | 24 | 24 |  | 23 | 25 | 25 | 142 |
| 7 | 18 | 22 | 21 |  | 19 | 20 | 19 | 119 |
| 8 | 20 | 25 | 25 |  | 23 | 24 | 23 | 140 |
| 9 | 21 | 24 | 25 |  | 25 | 25 | 26 | 146 |
| 10 | 22 | 25 | 25 |  | 23 | 24 | 22 | 141 |
| 11 | 21 | 22 | 23 |  | 24 | 24 | 23 | 137 |
| 12 | 20 | 21 | 20 |  | 20 | 22 | 22 | 125 |
| 13 | 23 | 25 | 24 |  | 21 | 23 | 21 | 137 |
| 14 | 24 | 26 | 25 |  | 22 | 21 | 21 | 139 |
| 15 | 23 | 23 | 23 |  | 25 | 25 | 24 | 143 |
| 16 | 19 | 24 | 23 |  | 24 | 25 | 25 | 140 |
| 17 | 19 | 20 | 21 |  | 22 | 23 | 23 | 128 |
| 18 | 23 | 23 | 22 |  | 23 | 23 | 23 | 137 |
| 19 | 22 | 26 | 26 |  | 20 | 19 | 19 | 132 |
| 20 | 22 | 25 | 22 |  | 24 | 26 | 20 | 139 |
| 21 | 18 | 21 | 21 |  | 18 | 19 | 19 | 116 |
| 22 | 21 | 25 | 23 |  | 23 | 22 | 24 | 138 |
| 23 | 19 | 23 | 21 |  | 19 | 22 | 20 | 124 |
| 24 | 22 | 25 | 25 |  | 23 | 24 | 23 | 142 |
| 25 | 24 | 27 | 27 |  | 20 | 23 | 23 | 144 |
| 26 | 22 | 24 | 25 |  | 23 | 23 | 24 | 141 |
| 27 | 23 | 26 | 27 |  | 21 | 24 | 24 | 145 |
| 28 | 24 | 26 | 25 |  | 25 | 20 | 23 | 143 |
