## Additional file 4 for "Microbiability of meat quality and carcass composition traits in swine"

Table S7. Variance components explained by microbiome relationship matrix (O), genomic relationship matrix (G), pen (P), residual (R), microbiability (m^2^) and heritability (h^2^) in different models^1^

| Traits^2^ | Effects | Model 0 | Model 1 | Model 2 | Model 3 |
| --- | --- | --- | --- | --- | --- |
| LD | P | 6.55±1.67 | 6.55±1.67 | 6.42±1.67 | 6.49±1.66 |
|  | O | - | 0.29E-04±0.00 | 1.76±1.58 | 0.71±1.05 |
|  | G | 6.15±2.94 | 6.15±2.93 | 5.88±2.36 | 5.88±2.36 |
|  | R | 38.53±2.06 | 38.53±2.06 | 37.20±2.15 | 38.09±2.05 |
|  | m^2^ | - | 0.00±0.00 | 0.03+0.03 | 0.01±0.02 |
|  | h^2^ | 0.12±0.04 | 0.11±0.04 | 0.12±0.04 | 0.11±0.05 |
| FD | P | 2.78±0.62 | 2.37±0.60 | 2.90±0.67 | 2.77±0.65 |
|  | O | - | 5.37±1.05 | 2.71±0.87 | 0.18±0.48 |
|  | G | 9.84±1.55 | 7.61±1.34 | 9.00±1.48 | 9.80±1.56 |
|  | R | 9.99±1.05 | 6.56±1.05 | 7.83±1.13 | 9.84±1.14 |
|  | m^2^ | - | 0.25±0.04 | 0.12+0.03 | 0.01±0.02 |
|  | h^2^ | 0.44±0.05 | 0.34±0.05 | 0.40±0.05 | 0.43±0.05 |
| CADG | P | 0.78E-05±0.00 | 0.59E-05±0.00 | 0.61E-05 ± 0.00 | 0.72E-0±0.00 |
|  | O | - | 1.18±0.29 | 0.95±0.26 | 0.31±0.18 |
|  | G | 1.05±0.23 | 0.94±0.27 | 1.07±0.29 | 0.98±0.08 |
|  | R | 4.71±0.29 | 3.18±0.30 | 3.31±0.26 | 4.63±0.30 |
|  | m^2^ | - | 0.22±0.05 | 0.18+0.04 | 0.06±0.03 |
|  | h^2^ | 0.20±0.05 | 0.18±0.04 | 0.20±0.05 | 0.18±0.05 |
| HAM | P | 0.87E-07±00 | 0.73E-05±0.0 | 0.72E-05±0.00 | 0.85E-05±0.00 |
|  | O | - | 0.83±0.29 | 0.80±0.25 | 0.10±0.16 |
|  | G | 0.72±0.27 | 0.73±0.27 | 0.82±0.28 | 0.70±0.27 |
|  | R | 4.71±0.29 | 3.96±0.33 | 3.96±0.33 | 4.65±0.32 |
|  | m^2^ | - | 0.15±0.05 | 0.14+0.04 | 0.02±0.02 |
|  | h^2^ | 0.13±0.05 | 0.13±0.04 | 0.13±0.05 | 0.13±0.04 |
| LOIN | P | 0.82E-05±0.0 | 0.70E-05±00 | 0.70E-05±00 | 0.80E-05±0.00 |
|  | O | - | 0.44±0.18 | 0.43±0.16 | 0.10±0.16 |
|  | G | 0.62±0.18 | 0.64±0.18 | 0.65±0.18 | 0.58±0.17 |
|  | R | 2.87±0.18 | 2.45±0.21 | 2.46±0.21 | 2.79±0.19 |
|  | m^2^ | - | 0.13±0.05 | 0.12+0.04 | 0.03±0.02 |
|  | h^2^ | 0.18±0.05 | 0.18±0.04 | 0.18±0.05 | 0.17±0.05 |
| BEL | P | 0.79E-05±0.0 | 0.54E-05±0.00 | 0.59E-05±0.00 | 0.76E-05±0.00 |
|  | O | - | 1.89±0.38 | 1.37±0.38 | 0.28±0.20 |
|  | G | 1.34±0.36 | 1.18±0.33 | 1.39±0.36 | 1.28±0.35 |
|  | R | 5.09±0.34 | 3.50±0.37 | 3.45±0.38 | 4.88±0.36 |
|  | m^2^ | - | 0.29±0.05 | 0.20+0.04 | 0.04±0.03 |
|  | h^2^ | 0.21±0.05 | 0.18±0.04 | 0.21±0.05 | 0.19±0.05 |
| IMF | P | 0.04 ± 0.02 | 0.04 ± 0.02 | 0.04± 0.02 | 0.04±0.02 |
|  | O | - | 0.06±0.03 | 0.03±0.02 | 0.03±0.02 |
|  | G | 0.55±0.08 | 0.53±0.08 | 0.54±0.08 | 0.55±0.08 |
|  | R | 0.41±0.05 | 0.37±0.37 | 0.39±0.05 | 0.38±0.04 |
|  | m^2^ | - | 0.06±0.02 | 0.03+0.02 | 0.03±0.02 |
|  | h^2^ | 0.55±0.05 | 0.53±0.05 | 0.54±0.05 | 0.54±0.05 |
| SMARB | P | 0.08 ± 0.02 | 0.08 ± 0.02 | 0.08 ± 0.02 | 0.08 ± 0.02 |
|  | O | - | 0.007±0.02 | 0.06±0.02 | 0.02±0.03 |
|  | G | 0.26±0.08 | 0.26±0.06 | 0.26±0.06 | 0.26±0.06 |
|  | R | 0.48±0.04 | 0.47±0.05 | 0.43±0.05 | 0.46±0.04 |
|  | m^2^ | - | 0.01±0.02 | 0.07+0.02 | 0.02±0.02 |
|  | h^2^ | 0.32±0.05 | 0.32±0.05 | 0.31±0.05 | 0.32±0.05 |
| MINA | P | 0.19 ± 0.04 | 0.17 ± 0.03 | 0.18 ± 0.03 | 0.18 ± 0.03 |
|  | O | - | 0.12±0.05 | 0.02±0.04 | 0.23E-04±0.00 |
|  | G | 0.22±0.06 | 0.22±0.06 | 0.22±0.06 | 0.22±0.06 |
|  | R | 0.85±0.06 | 0.77±0.07 | 0.83±0.05 | 0.85±0.06 |
|  | m^2^ | - | 0.09±0.02 | 0.02+0.02 | 0.00±0.00 |
|  | h^2^ | 0.17±0.05 | 0.16±0.05 | 0.17±0.05 | 0.17±0.05 |
| MINB | P | 0.15 ± 0.03 | 0.13 ± 0.03 | 0.14 ± 0.03 | 0.15 ± 0.03 |
|  | O | - | 0.08±0.03 | 0.005±0.01 | 0.64E-05±0.00 |
|  | G | 0.056±0.03 | 0.056±0.03 | 0.058±0.03 | 0.056±0.03 |
|  | R | 0.49±0.03 | 0.42±0.04 | 0.48±0.04 | 0.49±0.04 |
|  | m^2^ | - | 0.11±0.04 | 0.007+0.02 | 0.00±0.00 |
|  | h^2^ | 0.08±0.04 | 0.08±0.04 | 0.08±0.04 | 0.08±0.04 |
| MINL | P | 6.15 ± 1.16 | 6.15 ± 1.16 | 6.15 ± 1.16 | 6.15 ± 1.16 |
|  | O | - | 1.16±1.15 | 0.99E-05±0.00 | 0.61E-05±0.00 |
|  | G | 6.90±1.82 | 6.57±1.78 | 6.91±1.82 | 6.91±1.82 |
|  | R | 20.05±1.63 | 19.23±1.81 | 20.04±1.52 | 20.04±1.52 |
|  | m^2^ | - | 0.03±0.03 | 0.00+0.00 | 0.00±0.00 |
|  | h^2^ | 0.21±0.04 | 0.19±0.05 | 0.21±0.05 | 0.21±0.05 |
| PH | P | 0.013 ± 0.002 | 0.013 ± 0.002 | 0.013 ± 0.002 | 0.013 ± 0.002 |
|  | O | - | 0.002±0.002 | 0.001±0.007 | 0.26E-05±0.00 |
|  | G | 0.003±0.001 | 0.003±0.001 | 0.003±0.001 | 0.003±0.001 |
|  | R | 0.031±0.002 | 0.031±0.002 | 0.031±0.002 | 0.031±0.002 |
|  | m^2^ | - | 0.04±0.03 | 0.002+0.01 | 0.00±0.00 |
|  | h^2^ | 0.06±0.04 | 0.06±0.04 | 0.06±0.04 | 0.06±0.04 |
| SCOL | P | 0.014 ± 0.006 | 0.013 ± 0.006 | 0.014 ± 0.006 | 0.013 ± 0.006 |
|  | O | - | 0.012±0.011 | 0.40E-05±0.00 | 0.41E-05±0.00 |
|  | G | 0.096±0.02 | 0.097±0.02 | 0.096±0.02 | 0.096±0.02 |
|  | R | 0.216±0.018 | 0.204±0.019 | 0.215±0.018 | 0.215±0.018 |
|  | m^2^ | - | 0.04±0.03 | 0.002+0.01 | 0.00±0.00 |
|  | h^2^ | 0.30±0.06 | 0.30±0.06 | 0.30±0.06 | 0.30±0.05 |
| SFIRM | P | 0.026 ± 0.029 | 0.022 ± 0.028 | 0.028 ± 0.029 | 0.026 ± 0.029 |
|  | O | - | 0.012±0.011 | 0.40E-05±0.00 | 0.41E-05±0.00 |
|  | G | 0.134±0.050 | 0.122±0.046 | 0.120±0.044 | 0.134±0.050 |
|  | R | 0.904±0.059 | 0.791±0.068 | 0.834±0.063 | 0.904±0.059 |
|  | m^2^ | - | 0.13±0.04 | 0.08+0.03 | 0.00±0.00 |
|  | h^2^ | 0.13±0.04 | 0.11±0.04 | 0.11±0.04 | 0.13±0.05 |
| SSF | P | 0.52±0.36 | 0.49±0.36 | 0.50±0.36 | 0.52±0.36 |
|  | O | - | 0.30±0.36 | 1.38±0.59 | 0.41±0.32 |
|  | G | 3.07±0.74 | 3.03±0.73 | 3.00±0.73 | 3.12±0.75 |
|  | R | 9.42±0.70 | 9.20±0.66 | 8.27±0.78 | 8.99±0.77 |
|  | m^2^ | - | 0.02±0.02 | 0.10+0.04 | 0.03±0.02 |
|  | h^2^ | 0.24±0.05 | 0.23±0.05 | 0.22±0.05 | 0.24±0.05 |

^1^ Model 0 contains **G** matrix and pen effect as random effect, Model 1, Model 2 and Model 3 contains **O** matrix at weaning, mid test and off test in addition to **G** matrix and pen effect.

^2^LD = Loin depth; FD = Fat depth; CADG = Carcass average daily gain; IMF = Intramuscular fat percent, MINA = Minolta a*, MINB = Minolta b*, MINL = Minolta L*, PH = Ultimate pH; SCOL = Subjective color score; SMARB = Subjective marbling score; SFIRM = Subjective firmness score; SSF = Slice shear force, HAM = Ham weight; LOIN = Loin weight; BEL = Belly weight

Table S8: Variance components explained by microbiome relationship matrix (O), pen (P), residual (R) and microbiability (m^2^) at different stages of production when only microbiome information was included in the model

| Traits^2^ | Effects | Weaning | Mid test | Off test |
| --- | --- | --- | --- | --- |
| LD | P | 7.35±1.71 | 7.30±1.71 | 7.47±1.71 |
|  | O | 1.16±1.19 | 2.27±1.69 | 2.3E-05±0.00 |
|  | R | 42.52±2.35 | 41.62±2.48 | 43.56±2.17 |
|  | m^2^ | 0.03±0.02 | 0.04±0.03 | 0.00±0.00 |
| FD | P | 4.33±0.79 | 4.49±0.78 | 3.76±0.71 |
|  | O | 0.38±0.57 | 3.64±1.03 | 6.96±1.24 |
|  | R | 17.59±1.01 | 14.29±1.01 | 11.54±0.97 |
|  | m^2^ | 0.02±0.02 | 0.16±0.04 | 0.31±0.05 |
| CADG | P | 4.86E-06±0.00 | 4.86E-07±0.00 | 4.86E-07±0.00 |
|  | O | 0.36±0.19 | 0.97±0.27 | 1.20±0.3 |
|  | R | 4.80±0.25 | 4.27±0.26 | 4.03±0.27 |
|  | m^2^ | 0.07±0.03 | 0.19±0.04 | 0.23±0.09 |
| HAM | P | 2.09E-08±0.00 | 2.63E-07±0.00 | 2.57E-07±0.00 |
|  | O | 0.08±0.01 | 0.41±0.27 | 0.50±0.02 |
|  | R | 2.89±0.02 | 2.60±0.26 | 2.52±0.02 |
|  | m^2^ | 0.03±0.03 | 0.13±0.04 | 0.16±0.05 |
| LOIN | P | 4.24E-06±0.00 | 3.07E-07±0.00 | 3.03E-07±0.00 |
|  | O | 0.19±0.12 | 0.41±0.16 | 0.43±0.02 |
|  | R | 3.27±0.17 | 3.07±0.18 | 3.06±0.18 |
|  | m^2^ | 0.06±0.03 | 0.12±0.04 | 0.14±0.05 |
| BEL | P | 4.71E-06±0.00 | 3.95E-07±0.00 | 3.64E-07±0.00 |
|  | O | 0.29±0.12 | 1.18±0.35 | 1.47±0.39 |
|  | R | 4.69±0.17 | 3.95±0.31 | 3.56±0.32 |
|  | m^2^ | 0.06±0.03 | 0.22±0.04 | 0.29±0.05 |
| IMF | P | 0.12±0.03 | 0.10±0.03 | 0.11±0.03 |
|  | O | 0.04±0.03 | 0.05±0.03 | 0.10±0.04 |
|  | R | 0.80±0.05 | 0.79±0.05 | 0.74±0.05 |
|  | m^2^ | 0.04±0.03 | 0.05±0.03 | 0.11±0.04 |
| SMARB | P | 0.11±0.03 | 0.10±0.03 | 0.11±0.03 |
|  | O | 6E-08±0.00 | 0.05±0.03 | 0.02±0.02 |
|  | R | 0.69±0.03 | 0.64±0.04 | 0.67±0.04 |
|  | m^2^ | 0.00±0.00 | 0.07±0.03 | 0.03±0.02 |
| MINA | P | 0.25±0.05 | 0.24±0.04 | 0.22±0.03 |
|  | O | 9E-08±0.00 | 0.03±0.03 | 0.15±0.06 |
|  | R | 1.12±0.05 | 0.99±0.06 | 0.90±0.06 |
|  | m^2^ | 0.00±0.00 | 0.03±0.03 | 0.12±0.04 |
| MINB | P | 0.14±0.03 | 0.15±0.02 | 0.14±0.03 |
|  | O | 3.8E-06±0.00 | 0.007±0.01 | 0.08±0.03 |
|  | R | 0.54±0.05 | 0.53±0.03 | 0.47±0.03 |
|  | m^2^ | 0.00±0.00 | 0.01±0.02 | 0.11±0.04 |
| MINL | P | 6.53±1.12 | 6.93±1.23 | 6.66±1.21 |
|  | O | 2.4E-06±0.00 | 0.18±0.67 | 2.70±1.45 |
|  | R | 23.4±1.17 | 25.76±1.42 | 23.66±1.63 |
|  | m^2^ | 0.00±0.00 | 0.005±0.02 | 0.08±0.04 |
| PH | P | 0.012±0.002 | 0.013±0.002 | 0.013±0.002 |
|  | O | 1.5E-09±0.00 | 0.00016±0.0007 | 0.002±0.001 |
|  | R | 0.033±0.001 | 0.033±0.002 | 0.031±0.002 |
|  | m^2^ | 0.00±0.00 | 0.003±0.01 | 0.04±0.03 |
| SCOL | P | 0.03±0.01 | 0.03±0.01 | 0.03±0.00 |
|  | O | 1.2E-09±0.00 | 1.8E-07±0.00 | 0.03±0.01 |
|  | R | 0.29±0.01 | 0.29±0.01 | 0.280.01 |
|  | m^2^ | 0.00±0.00 | 0.00±0.00 | 0.06±0.04 |
| SFIRM | P | 0.05±0.03 | 0.05±0.03 | 0.04±0.03 |
|  | O | 4.2E-07±0.00 | 5.8E-07±0.04 | 0.14±0.05 |
|  | R | 1.00±0.05 | 0.97±0.05 | 0.88±0.05 |
|  | m^2^ | 0.00±0.00 | 0.00±0.00 | 0.14±0.04 |
| SSF | P | 1.34±0.42 | 1.28±0.41 | 1.29±0.42 |
|  | O | 0.23±0.36 | 1.63±0.64 | 0.38±0.05 |
|  | R | 11.55±0.68 | 10.40±0.71 | 11.43±0.67 |
|  | m^2^ | 0.01±0.02 | 0.12±0.05 | 0.03±0.03 |

^2^LD = Loin depth; FD = Fat depth; CADG = Carcass average daily gain; IMF = Intramuscular fat percent, MINA = Minolta a*, MINB = Minolta b*, MINL = Minolta L*, PH = Ultimate pH; SCOL = Subjective color score; SMARB = Subjective marbling score; SFIRM = Subjective firmness score; SSF = Slice shear force, HAM = Ham weight; LOIN = Loin weight; BEL = Belly weight
