## Additional file 5 for "Microbiability of meat quality and carcass composition traits in swine"

Table S9. Estimates of genomic correlation (below diagonal) at end of test among meat quality traits.

|  | **^1^SCOL** | **IMF** | **SFIRM** | **MINA** | **MINB** | **PH** |
| --- | --- | --- | --- | --- | --- | --- |
| **SCOL** |  |  |  |  |  |  |
| **IMF** | -0.24±0.13 |  |  |  |  |  |
| **SFIRM** | 0.16±0.19 | **^2^0.36±0.15** |  |  |  |  |
| **MINA** | **0.45±0.16** | **0.29±0.14** | -0.38±0.26 |  |  |  |
| **MINB** | **-0.94±0.22** | **0.78±0.16** | -0.06±0.31 | -0.02±010 |  |  |
| **PH** | **0.91±0.29** | -0.18±0.25 | 0.42±0.35 | -0.05±0.31 | -0.53±0.42 |  |

Table S10. Estimates of genomic correlation (below diagonal) at end of test among carcass composition traits.

|  | **^1^FD** | **CADG** | **HAM** | **LOIN** | **BEL** |
| --- | --- | --- | --- | --- | --- |
| **FD** |  |  |  |  |  |
| **CADG** | **0.27±0.13** |  |  |  |  |
| **HAM** | 0.03±0.17 | **0.67±0.13** |  |  |  |
| **LOIN** | -0.11±0.15 | **0.69±0.10** | **0.54±0.19** |  |  |
| **BEL** | **0.62±0.11** | **0.79±0.07** | **0.58±0.19** | **0.70±0.03** |  |

Table S11. Estimates of genomic correlation between meat quality traits and carcass composition traits with inclusion of microbiome

|  | **^1^FD** | **CADG** | **HAM** | **LOIN** | **BEL** |
| --- | --- | --- | --- | --- | --- |
| **SCOL** | 0.06±0.13 | -0.10±0.17 | -0.11±0.19 | -0.04±0.16 | -0.07±0.17 |
| **IMF** | **0.23±0.11** | 0.08±0.15 | -0.05±0.16 | -0.03±0.14 | **0.28±0.14** |
| **SFIRM** | **0.27±0.13** | 0.09±0.23 | 0.09±0.25 | 0.10±0.22 | **0.46±0.23** |
| **MINA** | **0.27±0.13** | -0.10±0.21 | -0.31±0.22 | -0.24±0.21 | -0.15±0.21 |
| **MINB** | **0.42±0.21** | 0.19±0.26 | 0.02±0.20 | 0.08±0.26 | 0.13±0.27 |
| **PH** | -0.02±0.26 | 0.10±0.32 | -0.19±0.28 | 0.09±0.20 | -0.30±0.33 |

Table S12. Estimates of genomic correlation between meat quality traits and carcass composition traits without inclusion of microbiome

|  | **^1^FD** | **CADG** | **HAM** | **LOIN** | **BEL** |
| --- | --- | --- | --- | --- | --- |
| **SCOL** | -0.06±013 | -0.01±0.17 | -0.05±0.18 | -0.02±0.17 | -0.10±0.17 |
| **IMF** | **0.26±0.10** | 0.13±0.14 | 0.01±0.16 | -0.02±0.14 | **0.37±0.14** |
| **SFIRM** | **0.28±0.08** | 0.13±0.28 | 0.12±0.24 | 0.02±0.22 | **0.51±0.18** |
| **MINA** | **0.30±0.14** | 0.09±0.17 | -0.27±0.21 | -0.28±0.20 | -0.05±0.21 |
| **MINB** | **0.46±0.19** | 0.14±0.27 | 0.02±0.25 | -0.02±0.27 | 0.16±0.27 |
| **PH** | 0.01±0.24 | 0.12±0.31 | -0.28±0.36 | 0.06±0.31 | -0.26±0.32 |

^1^FD = Fat depth; CADG = Carcass average daily gain; IMF = Intramuscular fat percent, MINA = Minolta a*, MINB = Minolta b*, MINL = Minolta L*, PH = Ultimate pH; SCOL = Subjective color score; SMARB = Subjective marbling score; SFIRM = Subjective firmness score; HAM = Ham weight; LOIN = Loin weight; BEL = Belly weight;

^2^Numbers in bold are significant.
